# Implanted Nerve Electrical Stimulation allows to Selectively Restore Hand and Forearm Movements in Patients with a Complete Tetraplegia

**DOI:** 10.1101/534362

**Authors:** Wafa Tigra, Christine Azevedo, Jacques Teissier, Anthony Gelis, Bertrand Coulet, Jean-Louis Divoux, David Guiraud

**Affiliations:** Wafa Tigra, PhD, is an independent researcher and was with MXM, Vallauris, France and INRIA, University of Montpellier, Montpellier, France; Christine Azevedo, PhD, is with INRIA, University of Montpellier, Montpellier, France; Jacques Teissier, MD, is with BeauSoleil Clinical Center, Montpellier, France; Anthony Gelis, MD, PhD, is with Propara Neurological Center, Montpellier, France; Bertrand Coulet, MD, PhD, is with CHU Lapeyronie, Montpellier, France; Jean-Louis Divoux, MSc, was with Axonic, Vallauris, France; David Guiraud, PhD, is with INRIA, University of Montpellier, Montpellier, France

## Abstract

**Background:** Functional Electrical Stimulation (FES) is used for decades in rehabilitation centers. For patients with a spinal cord injury (SCI), FES can prevent muscular atrophy, reduce spasticity and/or restore limb movements. To this last aim, FES external devices can be used, but elicit imprecise movements. FES implanted devices used neuromuscular stimulation and required implantation of one electrode for each site (muscle) to stimulate, an heavy surgical procedure and 10 times more electrical charges than nerve stimulation. Moreover, complications related to the numerous implanted components could appear over time. The purpose of this work is to evaluate whether a multi-contact nerve cuff electrode can selectively activate forearm and hand muscles and restore functional movements in patient with a complete tetraplegia.

**Methods:** A 12-contacts cuff electrode was designed to selectively activate, without damage it, multi fascicular peripheral nerve (*1*). Six subjects rwith at least one extensor or one flexor muscle stimulable by surface electrical stimulation, have been enrolled into the study. During the surgical procedure, radial or median nerve was stimulated through up to 35 different configurations of stimulation at an intensity up to 2.1 mA (25 Hz, pulse width 250 *µ*s, inter-pulse 100 *µ*s). Surface electrodes were positioned upon the forearm to record compound muscle action potentials elicited by nerve electrical stimulation. Recruitments curves and videos were recorded.

**Results:** The multicontact cuff electrode allowed us to selectively activate group of muscles to produce multiple, independent and functional hand and forearm movements in persons with tetraplegia. A surface electrical stimulation did not allow all those movements.

## Introduction

The incidence of SCIs in western Europe and the United States is estimated at 16 and 40 cases per million, respectively (*2*), with the high proportion of cervical injuries increasing progressively (*3*). SCIs can have devastating impacts on patients’ health, autonomy and quality of life. Technical aids (e.g. motorized wheelchair, orthosis, medical electric bed, transfer board, home automation …) allow to give back some independence to people with tetraplegia, but recovery of gripping movements is still felt as the priority (*4-8*). Indeed, most activities of daily living are performed via hand movements and restoration of forearm, hand and wrist active motricity would significantly increase their independence and quality of life. Without spinal cord repair, only partial solutions are available. For several decades, functional surgeries based on tendon–muscle transfers, and more recently nerve transfers (*9*), have been used to improve the functional potential of people with tetraplegia. During tendon–muscle transfers, the distal part of a tendon and its functional muscle, is detached from its natural insertion point and then fixed on a non-functional adjacent muscle in order to restore its initial function. For example, transfer of the biceps brachii to the triceps brachii can restore elbow active extension, the residual flexion of the elbow being provided by the other flexors (brachialis and brachioradialis). However, these 2 methods require sufficient muscles or nerves under voluntary control, and rehabilitation does not allow for systematic recovery of desired movements. Therefore, some people with tetraplegia are not able to undergo neurotendinous surgery. In that case, implanted or external FES can be an alternative to restore grip movements. One of the first applications of FES on hand muscles is reported by Backhouse et al. in 1954 (*10*). Used to determine the function of muscles, FES was then used to recover grip movements in patients with a high tetraplegia as early as 1963 (*11-13*). Nowaday, only few devices using FES to restore or improve grip functions are available. All of these devices use intramuscular, epimysial or surface electrodes (*14*) and so, need one electrode for each muscle to activate. However, the use of these devices is still very limited since they have limits in terms of acceptability, efficiency and benefit/limitation ratio. Furthermore, to achieve gripping movements, the higher the patient’s level of injury is, the greater the number of muscles to stimulate. Activate more than one muscle by electrode becomes relevant. Neural stimulation would have the advantage to stimulate more muscles via a single electrode (and so reduce implanted components), and will require less energy for muscle activation. Studies have demonstrated the feasibility of this approach (*15-18*), muscle that was selectively activated was the first muscle to branch distal to the cuff location. However, they combined multisite neuromuscular stimulation which makes this approach complex and therefore difficult to use in clinical routine.

The use of multipolar electrodes (poles ≥4) placed just above nerve bifurcations would make possible, selective activation of several fascicles from the same nerve. Indeed, upper limb fascicles would tend to anastomose and separate over a large part of their length, but would be somatotopically organized distally (*19, 20*). This selective activation could potentially activate different functions and/or muscles independently. Recruitment of agonist muscles via a single electrode may also be possible. Alternative activation of agonist muscles could reduce muscle fatigue and provide a better control of the desired movements.

In this study, we describe an approach based exclusively on neural stimulation to restore hand movements through a twelve contacts cuff electrode placed around upper-arm nerve (radial or median). Each of these contacts can be activated independently and so, current can be steered in many ways into the nerve. This technology could be used in combinaison with sus-lesional EMG signals or movements (*21*) to control the nerve stimulation. Thereby, individuals with tetraplegia, who can not benefit from a musculotendinous surgery, would be provided with a hand neuroprosthesis, usable in a clinical context, and allowing grasp movements recovery.

## Material and Methods

### Patient selection and surgery

6 patients with a C5 complete motor cervical injury were included in our experimental protocol. The protocol was approved by the local ethics committee (#ID-RCB: 2014-A01752-45) and experiments performed in accordance with the Declaration of Helsinki. Each patient signed an informed consent. The tests were performed during a scheduled surgery - a tendon transfer to recover elbow extension - avoiding a dedicated surgery. Moreover the experimental time slot was 30 min to let the anesthesia below 2 hours in total. Thus, a unique nerve was intra-operatively tested on each patient. Median or radial nerve was chosen depending on the surgical approach. The protocol was composed of 2 sessions: the first one consisted in a functional mapping of the targeted muscles’ groups namely thumb, fingers and wrist flexion/extension that leads to the inclusion of the patient. The second session consisted in the surgery itself.

After the exposure of the nerve about 5 cm around the elbow, one cuff electrode was placed around the targeted nerve and then gently sutured to avoid its displacement (section 6 of (*22*)).

### Electrodes

A 4 mm diameter, 2 cm length cuff electrode was used for radial nerves (3×3 contacts, Cortec GmbH, Freiburg, Germany) and a 6 mm diameter, 2 cm length cuff electrode was used for median nerves (3×4 contacts, Cortec GmbH, Freiburg, Germany, cf. figure 1). The cuff electrode contacts (2.79 × 0.79 mm2, 5.9 mm spacing between two longitudinal adjacent contacts) were 90/10 Pt/Ir made and embedded with silicone (Nusil).

**Figure 1:**
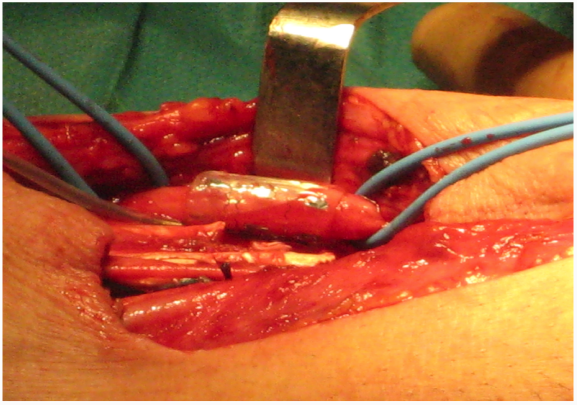
6 mm cuff electrode around the median nerve

### EMG and video recordings

Up to 3 pairs (bipolar recording) of sterile surface EMG electrodes were positioned on the forearm to capture group of muscles of interest (cf. table 2). The EMG were amplified with Gtec amplifier (ADinstrument, Austria) with the following settings: Amplification 1000, Low pass 1KHz, High pass 0.5Hz, 50Hz notch filter. The analog signal was sampled with LabChart (16 bits, Sampling Frequency: 10kHz) synchronized with the stimulator and a video (Logitech HD Pro Webcam C920 Refresh) capturing hand movements/surgeon’s comments (cf. fig. 2).

**Table 1:** Subject characteristics.

| Subject | Sex | Age (years) | Lesion time (months) | Nerve stimulated | Side operated | Giens<sup>1</sup> score |
| --- | --- | --- | --- | --- | --- | --- |
| P1 | M | 19 | 30 | median | right | 1 |
| P2 | M | 23 | 22 | radial | left | 0 |
| P3 | M | 25 | 34 | radial | left | 1 |
| P4 | M | 31 | 21 | radial | left | 2 |
| P5 | M | 32 | 29 | radial | left | 2 |
| P6 | M | 54 | 7 | median | right | 2 |

**Table 2:** Acronym of muscles of interest

| Acronym | Name |
| --- | --- |
| ECRL | Extensor Carpi Radialis Longus |
| ECRB | Extensor Carpi Radialis Brevis |
| EPL | Extensor Pollicis Longus |
| EDC | Extensor Digitorum Communis |
| FPL | Flexor Pollicis Longus |
| FDS/FDP | Flexor Digitorum Superficialis/Profundus |

**Figure 2:**
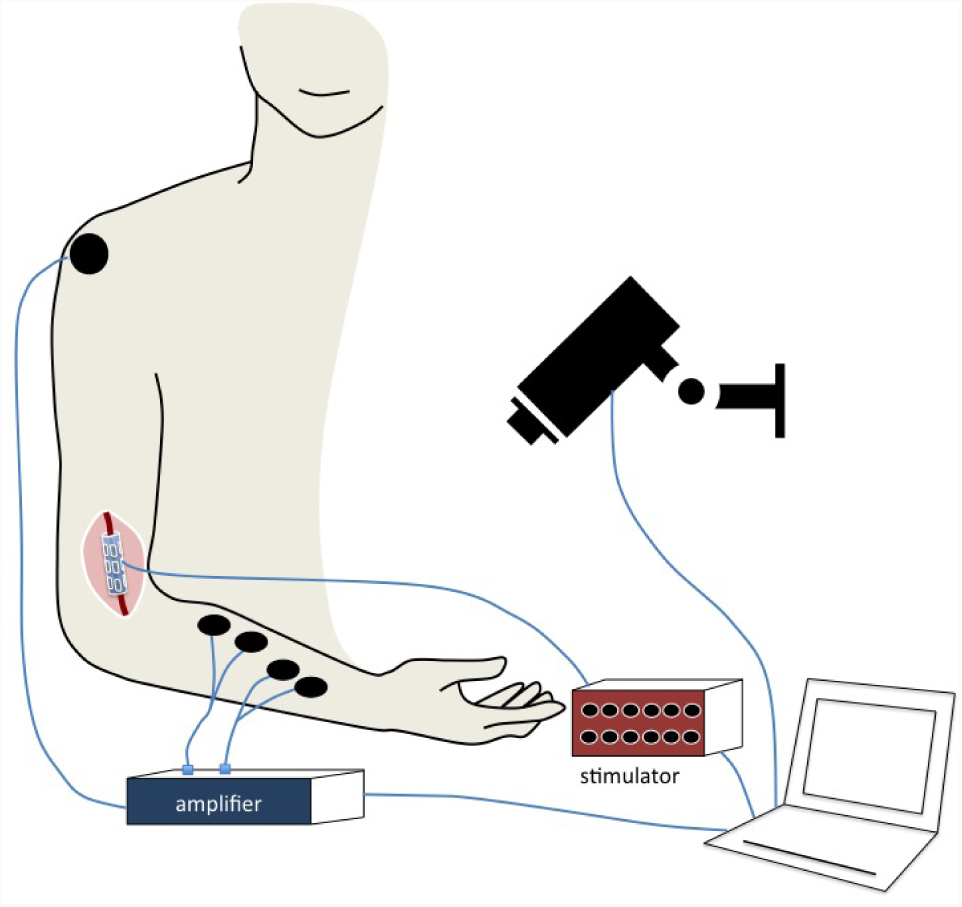
Setup design

### Stimulator

The stimulator was co-developed by Axonic and the University of Montpellier based on the architecture described in (*23*). The stimulator was able to deliver up to 5.1 mA, 20 *µ*A step, 0-511 *µ*s, 2 *µ*s step, 20 V compliance voltage. Moreover, the stimulator can distribute the current independently and in synchrony over the 12 contacts of the cuff electrode. The stimulator follows the essential requirements of safety concerning both the embedded software and hardware, with insulated interfaces. The waveform stimulation is rectangular, biphasic, asymmetric, charge balanced, and a delay between the cathodic and anodic phase allowing to decrease the quantity of charge needed without affecting considerably the selectivity of stimulation (*24*) is inserted.

### Software

The user can set the intensity (up to 2.4 mA), the pulse width (up to 600 *µ*s), the delay between the cathodic and anodic phase (up to 2 ms), the frequency (up to 50 Hz) and the distribution ratios (from 1/16 to 15/16) either cathodic or anodic.

### Stimulation protocol

To assess the selectivity of our multi-contact cuff electrode, we selected up to 35 configurations of stimulation (cf. tables 3 and 4) based on simulation studies (*25*) which showed that they provide a panel of shape of stimulation within the nerve that may fit the need for a selective fascicular activation.

**Table 3:** Configurations of stimulation sent for the median nerve about 5 cm from the elbow

| # | Name | Channel number |  |  |  |  |  |  |  |  |  |  |  |
| --- | --- | --- | --- | --- | --- | --- | --- | --- | --- | --- | --- | --- | --- |
|  |  | 1 | 2 | 3 | 4 | 5 | 6 | 7 | 8 | 9 | 10 | 11 | 12 |
| 1 | Ring | 2 | 1 | 2 | 1 | -3 | -3 | -3 | -3 | 2 | 1 | 2 | 1 |
| 2 | 1 Cathode<sub>B1</sub> 2 Anodes, Rings | 0 | 1 | 1 | 1 | -12 | 3 | 0 | 3 | 0 | 1 | 1 | 1 |
| 3 | 1 Cathode<sub>B2</sub> 2 Anodes, Rings | 1 | 0 | 1 | 1 | 3 | -12 | 3 | 0 | 1 | 0 | 1 | 1 |
| 4 | 1 Cathode<sub>B3</sub> 2 Anodes, Rings | 1 | 1 | 0 | 1 | 0 | 3 | -12 | 3 | 1 | 1 | 0 | 1 |
| 5 | 1 Cathode<sub>B4</sub> 2 Anodes, Rings | 1 | 1 | 1 | 0 | 3 | 0 | 3 | -12 | 1 | 1 | 1 | 0 |
| 6 | 1 Cathode<sub>B1</sub>, Rings | 1 | 2 | 1 | 2 | -12 | 0 | 0 | 0 | 1 | 2 | 1 | 2 |
| 7 | 1 Cathode<sub>B2</sub>, Rings | 2 | 1 | 2 | 1 | 0 | -12 | 0 | 0 | 2 | 1 | 2 | 1 |
| 8 | 1 Cathode<sub>B3</sub>, Rings | 1 | 2 | 1 | 2 | 0 | 0 | -12 | 0 | 1 | 2 | 1 | 2 |
| 9 | 1 Cathode<sub>B4</sub>, Rings | 2 | 1 | 2 | 1 | 0 | 0 | 0 | -12 | 2 | 1 | 2 | 1 |
| 10 | Longitudinal Tripolar<sub>B1</sub> | 6 | 0 | 0 | 0 | -12 | 0 | 0 | 0 | 6 | 0 | 0 | 0 |
| 11 | Longitudinal Tripolar<sub>B2</sub> | 0 | 6 | 0 | 0 | 0 | -12 | 0 | 0 | 0 | 6 | 0 | 0 |
| 12 | Longitudinal Tripolar<sub>B3</sub> | 0 | 0 | 6 | 0 | 0 | 0 | -12 | 0 | 0 | 0 | 6 | 0 |
| 13 | Longitudinal Tripolar<sub>B4</sub> | 0 | 0 | 0 | 6 | 0 | 0 | 0 | -12 | 0 | 0 | 0 | 6 |
| 14 | Bipolar<sub>AC</sub> | -3 | -3 | -3 | -3 | 0 | 0 | 0 | 0 | 3 | 3 | 3 | 3 |
| 15 | Bipolar<sub>CA</sub> | 3 | 3 | 3 | 3 | 0 | 0 | 0 | 0 | -3 | -3 | -3 | -3 |
| 16 | Bipolar<sub>BA</sub> | 3 | 3 | 3 | 3 | -3 | -3 | -3 | -3 | 0 | 0 | 0 | 0 |
| 17 | Bipolar<sub>AB</sub> | -3 | -3 | -3 | -3 | 3 | 3 | 3 | 3 | 0 | 0 | 0 | 0 |
| 18 | Bipolar<sub>BC</sub> | 0 | 0 | 0 | 0 | -3 | -3 | -3 | -3 | 3 | 3 | 3 | 3 |
| 19 | Bipolar<sub>CB</sub> | 0 | 0 | 0 | 0 | 3 | 3 | 3 | 3 | -3 | -3 | -3 | -3 |
| 20 | Steering<sub>B1</sub> | 0 | 1 | 1 | 1 | -12 | 0 | 6 | 0 | 0 | 1 | 1 | 1 |
| 21 | Steering<sub>B2</sub> | 1 | 0 | 1 | 1 | 0 | -12 | 0 | 6 | 1 | 0 | 1 | 1 |
| 22 | Steering<sub>B3</sub> | 1 | 1 | 0 | 1 | 6 | 0 | -12 | 0 | 1 | 1 | 0 | 1 |
| 23 | Steering<sub>B4</sub> | 1 | 1 | 1 | 0 | 0 | 6 | 0 | -12 | 1 | 1 | 1 | 0 |
| 1' | Ring | 2 | 1 | 2 | 1 | -3 | -3 | -3 | -3 | 2 | 1 | 2 | 1 |
| 24 | Transverse Tripolar<sub>A1</sub> | -14 | 7 | 0 | 7 | 0 | 0 | 0 | 0 | 0 | 0 | 0 | 0 |
| 25 | Transverse Tripolar<sub>A2</sub> | 7 | -14 | 7 | 0 | 0 | 0 | 0 | 0 | 0 | 0 | 0 | 0 |
| 26 | Transverse Tripolar<sub>A3</sub> | 0 | 7 | -14 | 7 | 0 | 0 | 0 | 0 | 0 | 0 | 0 | 0 |
| 27 | Transverse Tripolar<sub>A4</sub> | 0 | 0 | 0 | -14 | 7 | 0 | 7 | 0 | 0 | 0 | 0 | 0 |
| 28 | Transverse Tripolar<sub>B1</sub> | 0 | 0 | 0 | 0 | -14 | 7 | 0 | 7 | 0 | 0 | 0 | 0 |
| 29 | Transverse Tripolar<sub>B2</sub> | 0 | 0 | 0 | 0 | 7 | -14 | 7 | 0 | 0 | 0 | 0 | 0 |
| 30 | Transverse Tripolar<sub>B3</sub> | 0 | 0 | 0 | 0 | 0 | 7 | -14 | 7 | 0 | 0 | 0 | 0 |
| 31 | Transverse Tripolar<sub>B4</sub> | 0 | 0 | 0 | 0 | 7 | 0 | 7 | -14 | 0 | 0 | 0 | 0 |
| 32 | Transverse Tripolar<sub>C1</sub> | 0 | 0 | 0 | 0 | 0 | 0 | 0 | 0 | -14 | 7 | 0 | 7 |
| 33 | Transverse Tripolar<sub>C2</sub> | 0 | 0 | 0 | 0 | 0 | 0 | 0 | 0 | 7 | -14 | 7 | 0 |
| 34 | Transverse Tripolar<sub>C3</sub> | 0 | 0 | 0 | 0 | 0 | 0 | 0 | 0 | 0 | 7 | -14 | 7 |
| 35 | Transverse Tripolar<sub>C4</sub> | 0 | 0 | 0 | 0 | 0 | 0 | 0 | 0 | 7 | 0 | 7 | -14 |

**Table 4:** Configurations of stimulation sent for the radial nerve about 5 cm from the elbow

| # | Name | Channel number |  |  |  |  |  |  |  |  |
| --- | --- | --- | --- | --- | --- | --- | --- | --- | --- | --- |
|  |  | 1 | 2 | 3 | 4 | 5 | 6 | 7 | 8 | 9 |
| 1 bis | Ring | 2 | 2 | 2 | -4 | -4 | -4 | 2 | 2 | 2 |
| 2 bis | 1 Cathode<sub>B1</sub> 2 Anodes, Rings | 1 | 1 | 1 | -12 | 3 | 3 | 1 | 1 | 1 |
| 3 bis | 1 Cathode<sub>B2</sub> 2 Anodes, Rings | 1 | 1 | 1 | 3 | -12 | 3 | 1 | 1 | 1 |
| 4 bis | 1 Cathode<sub>B3</sub> 2 Anodes, Rings | 1 | 1 | 1 | 3 | 3 | -12 | 1 | 1 | 1 |
| 6 bis | 1 Cathode<sub>B1</sub>, Ring | 2 | 2 | 2 | -12 | 0 | 0 | 2 | 2 | 2 |
| 7 bis | 1 Cathode<sub>B2</sub>, Ring Anode | 2 | 2 | 2 | 0 | -12 | 0 | 2 | 2 | 2 |
| 8 bis | 1 Cathode<sub>B3</sub>, Ring Anode | 2 | 2 | 2 | 0 | 0 | -12 | 2 | 2 | 2 |
| 10 bis | Longitudinal Tripolar<sub>B1</sub> | 6 | 0 | 0 | -12 | 0 | 0 | 6 | 0 | 0 |
| 11 bis | Longitudinal Tripolar<sub>B2</sub> | 0 | 6 | 0 | 0 | -12 | 0 | 0 | 6 | 0 |
| 12 bis | Longitudinal Tripolar<sub>B3</sub> | 0 | 0 | 6 | 0 | 0 | -12 | 0 | 0 | 6 |
| 14 bis | Bipolar<sub>AC</sub> | -4 | -4 | -4 | 0 | 0 | 0 | 4 | 4 | 4 |
| 17 bis | Bipolar<sub>AB</sub> | -4 | -4 | -4 | 4 | 4 | 4 | 0 | 0 | 0 |
| 18 bis | Bipolar<sub>BC</sub> | 0 | 0 | 0 | -4 | -4 | -4 | 4 | 4 | 4 |
| 19 bis | Bipolar<sub>CB</sub> | 0 | 0 | 0 | 4 | 4 | 4 | -4 | -4 | -4 |
| 1' bis | Ring | 2 | 2 | 2 | -4 | -4 | -4 | 2 | 2 | 2 |
| 24 bis | Transverse Tripolar<sub>A1</sub> | -14 | 7 | 7 | 0 | 0 | 0 | 0 | 0 | 0 |
| 25 bis | Transverse Tripolar<sub>A2</sub> | 7 | -14 | 7 | 0 | 0 | 0 | 0 | 0 | 0 |
| 26 bis | Transverse Tripolar<sub>A3</sub> | 7 | 7 | -14 | 0 | 0 | 0 | 0 | 0 | 0 |
| 28 bis | Transverse Tripolar<sub>B1</sub> | 0 | 0 | 0 | -14 | 7 | 7 | 0 | 0 | 0 |
| 29 bis | Transverse Tripolar<sub>B2</sub> | 0 | 0 | 0 | 7 | -14 | 7 | 0 | 0 | 0 |
| 30 bis | Transverse Tripolar<sub>B3</sub> | 0 | 0 | 0 | 7 | 7 | -14 | 0 | 0 | 0 |
| 32 bis | Transverse Tripolar<sub>C1</sub> | 0 | 0 | 0 | 0 | 0 | 0 | -14 | 7 | 7 |
| 33 bis | Transverse Tripolar<sub>C2</sub> | 0 | 0 | 0 | 0 | 0 | 0 | 7 | -14 | 7 |
| 34 bis | Transverse Tripolar<sub>C3</sub> | 0 | 0 | 0 | 0 | 0 | 0 | 7 | 7 | -14 |

During the surgical procedure, we first searched for the intensity threshold of activation using the classical ring configuration (configuration #1 in table 3 and #1bis in table 4). Then an automatic scanning was programmed from approximately 80% to 250% of this threshold value. The pulse width and the frequency were fixed (25 Hz, 250 *µ*s), the configurations and the intensities were scanned every 2 seconds (1s ON-1s OFF) to avoid fatigue. The ring configuration was scanned first and last to check the stability of the response during the whole trial. During the scan the surgeon labelled the movements elicited and their quotation (on the MRC scale). A double check was performed off-line with the synchronous audio/video.

## Results

We studied the capacity of a 12-contacts nerve cuff electrode to selectively activate upper arm muscles. We have shown that it is possible to activate selectively, *via* current steering, close and distinct muscles innervated by the same upper limb nerve in patients with tetraplegia. No adverse effects were reported after the surgery. In particular, medial and radial nerve stimulation in six persons with tetraplegia allowed to **selectively** activate (muscular contraction scored at 3 or more on the MRC scale), muscles allowing (cf. supplementary videos) a:

- for patient I: wrist flexion,
- for patient II: thumb extension,
- for patient III: supination, wrist extension,
- for patient IV: wrist extension, thumb extension, fingers extension,
- for patient V: supination, elbow flexion, wrist extension,
- for patient VI: fingers flexion, thumb flexion, thumb opposition (key and palmar grips were also observed).

### Radial nerve stimulation

Tables 5 and 6 synthesize observations reported by the surgeon during the intervention and completed by the video analysis.

**Table 5:** Visual observation of selective movements [elicited with configuration # in table 4] when radial nerve was stimulated, robust configurations (#) are in **bold**. *Electrode turned at 180

| Patient ID | Selective movements<br>[elicited with configuration #] | Current amplitude, (µA) | Maximal MRC<sup>2</sup><br>quotation |
| --- | --- | --- | --- |
| 2 | Thumb extension [3, 17, 18 bis]<br>Finger extension [1' bis] | 188, 300, 300<br>375 | 3<br>2 |
| 3 | Wrist extension [1, 6, 7, 10, **11, 12, 14, 17, 18** bis]<br>Supination [3 bis]<br>Elbow flexion [1' bis] | 270, 150, 150, 180, 150, 540, 150, 150<br>300<br>330 | 4+<br>3<br>2 |
| 4 | Thumb extension [2, 28*, 32, 33, 34 bis]<br>Wrist extension [1, 6, 7, 8, 10, **11, 12, 14, 17, 18, 24, 25\***, 28, **33\*** bis]<br>Fingers extension [**32\*** bis] | 750, 1313, 963, 1663, 1094<br>525, 450, 525, 450, 525, 525, 525, 600, 525, 1094, 1313, 1838, 963<br>963 | 4+<br>5<br>3 |
| 5 | Elbow flexion [4, 28*, 34 bis]<br>Wrist extension [1, 8, 12, **14, 17, 24** bis]<br>Supination [1', **2, 3, 6, 7, 11, 18, 19, 28, 29, 30, 32\***, **34\*** bis] | 488, 1444, 919<br>375, 525, 525, 525, 450, 1313<br>375, 450, 525, 300, 300, 413, 413, 450, 656, 656, 1181, 919, 1313 | 3<br>4<br>3 |

**Table 6:**
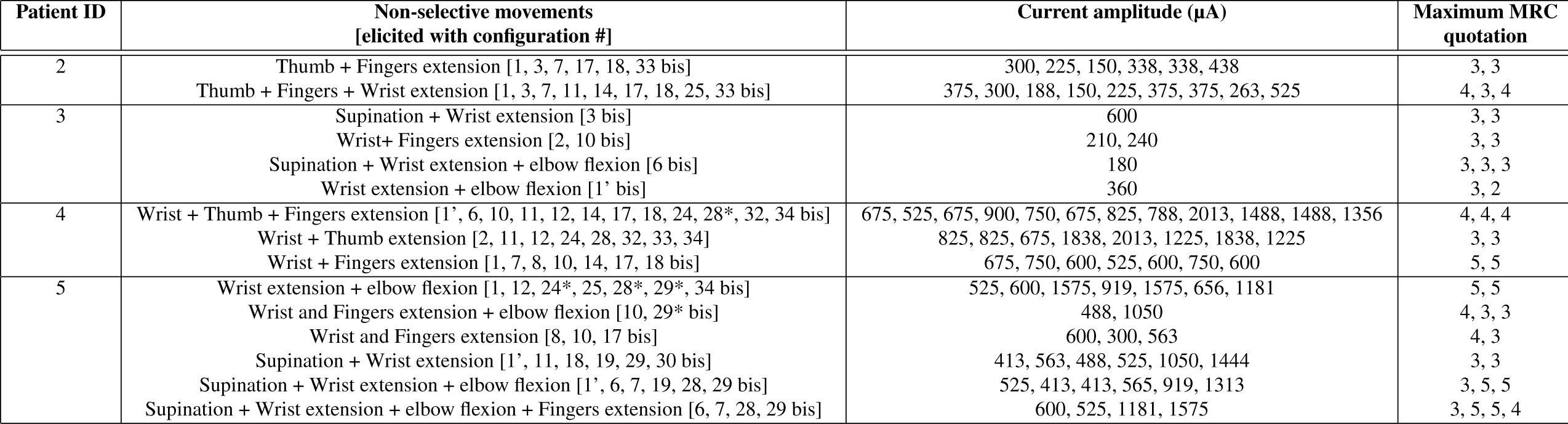
Visual observation of non-selective movements [elicited with configuration # in table 4] when radial nerve was stimulated, *Electrode turned at 180

### Recruitment order when the nerve is fully stimulated (Ring Configuration)

A full nerve stimulation (Ring) allows to selectively recrut the wrist extensors in 3 of 4 patients.

### Achieving selectivity with more than one configuration of stimulation

It is possible to produce the same selective movement with different configurations of stimulation (cf. table 5).

### Comparison of the “Ring” configuration repeated: significant fatigue

To examine the reproducibility of the results, we have repeated the “Ring” configuration of stimulation twice (at the beginning and end of the stimulation protocol). For 3 patients (P3, P4, P5) on 4, we were able to activate one more muscles when the configuration was repeated at the end of the scan and for 1 patient (P2) on 4, one less. In all cases, the MRC score decreased significantly, highlighting a fast muscular fatigue. The fatigue was also seen on the EMG signal amplitude.

### The same configuration of stimulation repeated at a distance of 5 mm and 10 mm does not activate the same muscles

The Transverse Tripolar configuration was repeated on each of the 12 contacts of the cuff electrode. For patient 4, a best selectivity was reached for the more distal ring (the one situated near the nerve bifurcation). For patient 5, we activate different muscles when the stimulation was repeated at 5 mm and 10 mm distance.

### Robustness of configurations

We consider a selective configuration as robust when an increase of 50% of the minimal intensity required to activate the muscle, does not affect the selectivity. For many configurations, a small increase of intensity activated other muscles (low robustness). Table 5 shows selective and robust configurations, for patients 3, 4 and 5

- Patient 3: Ring; 1 Cathode_*B*2_ 2 Anodes, Rings; 1 Cathode_*B*2_, Ring Anode; Longitudinal Tripolar_*B*2_; Longitudinal Tripolar_*B*3_; Bipolar_*AC*_; Bipolar_*AB*_; Bipolar_*BC*_. For this patient, the Tripolar Transverse configurations were not launched because of the very high muscular fatigue seen after few minutes of nerve stimulation.
- Patient 4: Ring; Longitudinal Tripolar_*B*2_; Transverse Tripolar_*A*1_; Transverse Tripolar_*A*2_; Transverse Tripolar_*C*1_; Transverse Tripolar_*C*2_
- Patient 5: 1 Cathode_*B*1_ 2 Anodes, Rings; 1 Cathode_*B*2_ 2 Anodes, Rings; 1 Cathode_*B*3_ 2 Anodes, Rings; Bipolar_*AC*_; Transverse Tripolar_*A*1_; Transverse Tripolar_*C*1_; Transverse Tripolar_*C*3_

### Median nerve stimulation

Tables 7 and 8 synthesize observations reported by the surgeon during the intervention and completed by the video analysis.

**Table 7:**
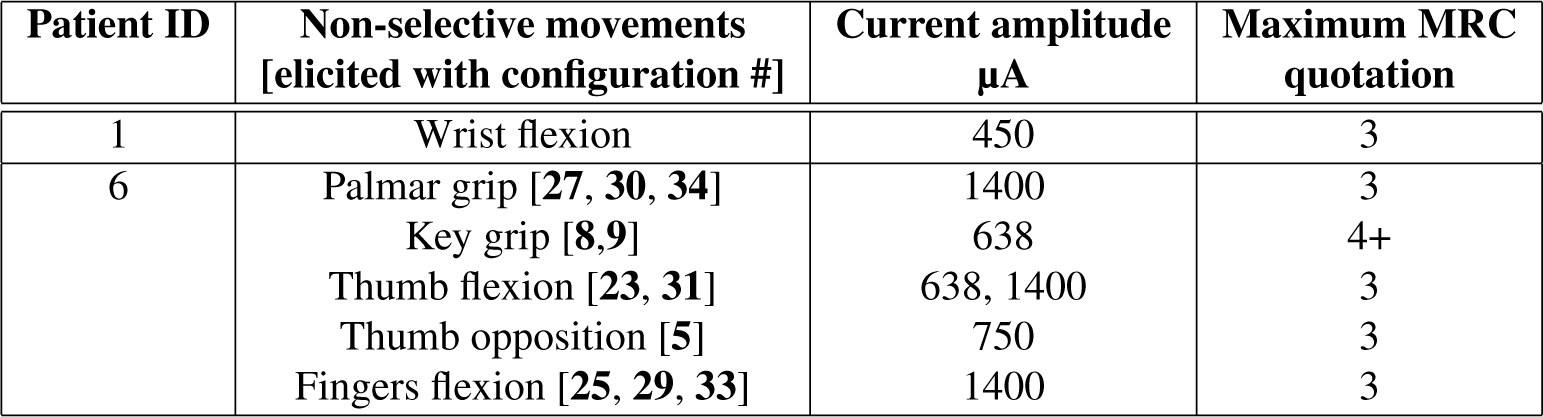
Visual observation of selective movements [elicited with configuration # in table 3] when median nerve was stimulated, robust configurations (#) are in **bold**

| Patient ID | Non-selective movements<br>[elicited with configuration #] | Current amplitude<br>$\mu\text{A}$ | Maximum MRC<br>quotation |
| --- | --- | --- | --- |
| 1 | Wrist flexion | 450 | 3 |
| 6 | Palmar grip [**27, 30, 34**] | 1400 | 3 |
|  | Key grip [**8,9**] | 638 | 4+ |
|  | Thumb flexion [**23, 31**] | 638, 1400 | 3 |
|  | Thumb opposition [**5**] | 750 | 3 |
|  | Fingers flexion [**25, 29, 33**] | 1400 | 3 |

**Table 8:** Visual observation of non-selective movements [elicited with configuration # in table 3] when median nerve was stimulated

| Patient ID | Non-selective movements<br>[elicited with configuration #] | Current amplitude<br>$\mu\text{A}$ | Maximum MRC<br>quotation |
| --- | --- | --- | --- |
| 6 | Thumb, Fingers flexion [24] | 1400 | 3, 3 |
|  | Thumb and Fingers flexion + thumb opposition [20, 28] | 638, 1400 | 3, 3 |
|  | Wrist, Fingers and Thumb flexion [1, 2, 3, 4, 6, 7, 8, 9, 10, 11, 12, 13, 14, 16, 17, 18, 21] | 638 | 4, 3, 3 |

### Recruitment order when the nerve is fully stimulated (Ring Configuration)

A full nerve stimulation (Ring) allows to activate a wrist, fingers and thumb flexion in patient 6.

### Achieving selectivity with more than one configuration of stimulation

It is possible to produce the same selective movement with different configurations of stimulation (cf. table 7).

### Comparison of the “Ring” configuration repeated: significant fatigue

To examine the reproducibility of the results, we have repeated the “Ring” configuration of stimulation twice (at the beginning and end of the stimulation protocol). We obtain the same results for patient 6, yet with an MRC score decreased significantly, highlighting a fast fatigue. The fatigue was also seen on the EMG signal amplitude.

### The same configuration of stimulation repeated at 5 mm and 10 mm distance does not activate the same muscles

The Transverse Tripolar configuration was repeated on each contact of the 12 contacts cuff electrode. For patient 6, we recruited more muscles when we stimulated the nerve near the bifurcation.

### Robustness of configurations

We consider a selective configuration as robust when an increase of 50% of the minimal intensity required to activate the muscle does not affect the selectivity. For many configurations, a small increase of intensity activated other muscles. Table 7 shows selective and robust configurations for patient 6.

## Discussion

Individuals with tetraplegia can decrease their use of human help and increase their functional independence thanks to neuroprostheses, particularly in activities of daily living requiring hand movements. Systems using muscular surface electrodes are bigger, require more energy to stimulate muscles and are less selective and reliable than systems using implanted electrodes. Currently, there is no commercialized implanted neuroprosthesis available to restore hand movements in persons with tetraplegia. The functional improvement provided by the Freehand system was considerable and impossible to achieve by other means available. However, this system had shown several disadvantages (*13, 26–29*). One can cite the numerous implanted materials lengthening the duration of the surgery and limiting drastically the possibility of bilateralization, essential to some activities of daily living. Human assistance needed daily for placement of external components and poor ergonomic control mode were additional limits. In this study, we have shown that it is possible, thanks to current steering allowed by a 12-contacts cuff electrode, to selectively activate, close and distinct muscles innervated by the same upper limb nerve. Moreover, some of the superficial muscles were activated during the second step (implanted nerve stimulation) but were not during the first step (surface muscle stimulation) performed before the surgery.

In the past, use of multipolar cuff electrodes, has selectively activated a single muscle, typically the first muscle to branch distal to the cuff location (*30*). In our study, the same pattern of stimulation reproduced at 0.5 and 1 cm distant led to distinct muscle recruitment patterns. This allows us to support that upper limb axons are reorganized within the same fascicle even near the nerve bifurcations or/and that some fascicles change their orientation.

Muscular atrophy and spasticity (excessif muscle contraction) are often seen in patient with a CNS injury and have been seen in patients involved in the study. Moreover, the same pattern reproduced at the beginning and the end of the scan showed a significant fatigue even though the nerve was stimulated for only few minutes. For these two last points, we can expect a functional improvement over time, if a muscle reinforcement by surface electrical stimulation takes place before the surgery.

During a nerve electrical stimulation, motor neurons with larger diameter (e.g. those that innervate type IIb muscle fibers, fast and fatigable) are activated first, followed by activation of smaller diameter motor neurons. This recruitment property is the opposite of the physiological recruitment order (*31*). Furthermore, the intensity of the stimulus produced by an electrical stimulation decreases with its distance from the source of stimulation. Therefore, the motor neurons closest to the electrode and of larger diameter are the most stimulated. Finally, contrary to a voluntary contraction, an electrical stimulation (neural or neuromuscular) recruits the muscle fibers in the same order. There is no longer a “rotation” system (*32*) for fibers recruitment and so, induces early muscle fatigue. In this study, we have seen that it is possible to activate the same muscles with different configurations of stimulation and so it would be possible to overcome some issues of electrical stimulation.

During our experiments, a weak space between the electrode and the nerve was present. We believe that the diameter of the cuff electrodes must be able to fit the exact diameter of the nerve. This, would optimally stimulate the nerve and allow ENG signals recording. These signals might be integrated into the neuroprosthesis and used as a closed-loop control.

## Conclusion

This study showed that two electrodes placed around two upper limb nerves allow to restore hand and forearm movements in patients with tetraplegia. Possible ways to pilot this stimulation have been investigated in previous studies (*33*), which showed that sus-lesional muscles can be used to pilot a surface electrical stimulation (unpublished data). Based on functional specifications revealed in these studies, we have demonstrated that a fully implantable system for restoring grip movements, with limited number of components could be developed. The technical simplicity of our approach would allow a bi-lateralization of the system, essential for some activities of daily living. Thus, a device using such a technology could, in combination with transfer surgery, be materially less heavy than those that existed previously. It would reduce the time required for the surgical procedure, risks related to the surgery, implanted materials and energy needed for its operation while increasing ergonomics. A balance between technical complexity and ease of use would be found. As feasibility has been demonstrated, a chronic clinical trial should be conducted shortly.

## Acknowledgments

The authors wish to thank the subjects involved into this research, MXM-Axonic/ANRT for support with the PhD grant (CIFRE # 2013/0867), Pawel Maciejasz, Chloé Picq, and Violaine Leynaert for valuable guidance and support.

## Footnotes

1 Giens classification is based on the number of remaining active muscles under the elbow in person with tetraplegia.

2 The MRC (Medical Research Council) scale allows to manually quantify the strength developed by each muscle. 0 corresponds to complete paralysis of the muscle, 1 to a minimal contraction, 2 to an active movement without gravity, 3 to a weak contraction against gravity, 4 to an active movement against gravity and resistance and 5 to a normal force.

